# Emergent topological structure in spontaneous brain-organoid activity

**DOI:** 10.64898/2026.07.17.739228

**Authors:** Eve Bodnia, Margaux Basart, Sofie Hai, Lenzie Ford, Nina Miolane, Kenneth S. Kosik, Dirk Bouwmeester, Lincoln D. Carr

**Affiliations:** Department of Molecular, Cellular, and Developmental Biology, University of California, Santa Barbara, CA 93106, USA; Department of Physics, University of California, Santa Barbara, CA 93106, USA; Department of Physics, Colorado School of Mines, Golden, CO 80401, USA; Quantum Engineering Program, Colorado School of Mines, Golden, CO 80401, USA; Department of Electrical and Computer Engineering, University of California, Santa Barbara, CA 93106, USA; Neuroscience Research Institute, University of California, Santa Barbara, CA 93106, USA; Huygens-Kamerlingh Onnes Laboratory, Leiden University, P.O. Box 9504, 2300 RA Leiden, the Netherlands; Department of Applied Mathematics and Statistics, Colorado School of Mines, Golden, CO 80401, USA; Institute of Science and Technology Austria (ISTA), Am Campus 1, 3400 Klosterneuburg, Austria

**Author notes:** These authors contributed equally to this work (Eve Bodnia, Margaux Basart).

## Abstract

Neural activity is widely held to organize on low-dimensional structure embedded in a high-dimensional state space. Persistent homology reads such structure directly from the pattern of pairwise correlations, without assuming in advance which variables are relevant. We apply persistent homology to microelectrode-array (MEA) recordings of spontaneous activity from human (Lancaster) and mouse (Paşca) cortical organoids, spanning 25–197 sorted units, and ask whether topological data analysis resolves structure at the node counts that neural recordings actually deliver. Building weighted networks in correlation space and characterizing them by Vietoris-Rips filtration, we find that the first homology (*H*_1_, loops) rises significantly above a rate- and population-preserving null in 14 of 18 datasets, all of which remain significant after correction for testing 18 datasets. Random removal of neural units leaves the loops largely intact. Removing the units that form the loops disrupts them more than removing equally connected units does. Topological richness grows with network size, and second homology (*H*_2_) exceeds the null only in the larger networks. These results show that persistent homology resolves structured topology in neural recordings at the scale experiments actually deliver.

## 1. Introduction

A recurring idea in systems neuroscience is that meaningful neural activity does not fill its high-dimensional state space but concentrates on a lower-dimensional manifold, the neural manifold hypothesis [1, 2, 3]. Where that manifold is curved or multiply connected, linear tools such as principal component analysis report the manifold’s dimension but miss its shape. Topological data analysis (TDA), and persistent homology in particular, recovers that shape, in the form of topological structure: the number and persistence of connected components, loops, and voids, counted respectively by the Betti numbers *β*_0_, *β*_1_, *β*_2_. These are computed directly from a matrix of pairwise relationships and are invariant to any monotone rescaling of that matrix [4, 5].

Applied to neural data, persistent homology has revealed the circular and toroidal geometry of spatial coding [2], intrinsic structure in hippocampal correlations independent of receptive-field models[4], cavities and cliques in structural connectomes and reconstructed microcircuits [6, 7], and homological organization of functional networks [8, 5]. In parallel, physicists studying complex systems have adopted higher-order interactions and simplicial structure as a common language [9, 10, 11]. The clique complex of a correlation matrix is built from pairwise relations, so its higher-dimensional homology describes how pairwise co-firing closes into loops and voids, not interactions among three or more units [9].

Brain organoids, self-organizing three-dimensional cultures of human or mouse cortical tissue [12, 13], are an increasingly important model system. High-density MEAs can now record the spontaneous spiking of hundreds of units in a single organoid at once [14]. Graph-theoretic network analysis has already been brought to organoid MEA data [15], characterizing degree, clustering, and modularity. Persistent homology characterizes the global organization of that same correlation graph, the loops and voids that local measures of this kind miss, and has not previously been applied to the spontaneous activity of brain organoids.

Real neural recordings, in organoids, in slices, and in vivo, rarely resolve more than a few hundred units at once, far short of the 10^4^ – 10^6^ in a full circuit. MEA is our starting point because its sub-millisecond sampling captures exact spike times, and the correlations we analyze are built on that timing; optical methods are one to two orders of magnitude slower and blur it. Across eighteen organoid datasets, persistent homology finds loop topology over and above what a rate- and population-preserving null produces, at the unit counts experiments actually deliver.

## 2. Organoid MEA data

We analyze spontaneous extracellular recordings from two organoid protocols: human cerebral organoids (Lancaster protocol; datasets labeled O*n*) and mouse cortical organoids (Paşca protocol; datasets labeled MO*n*). The protocols differ in ways that shape the networks: Lancaster organoids self-organize without regional patterning and develop numerous rosettes spanning several presumptive brain regions, whereas Paşca organoids are directed toward a cortical fate and carry one or few rosettes [12, 13]. Activity was recorded on high-density CMOS MEAs at 20 kHz for 3 min (O1 – O4), 10 – 12.7 min (MO1 – MO12), or 30 min (O5, O6), resolving spike times to sub-millisecond precision, and spike-sorted with Kilosort2 [16]. The sorted single units that passed quality control and the duplicate check below form the nodes of all networks. The resulting datasets span *N* = 25 to *N* = 197 units (Table 1).

**Table 1.** Datasets, unit counts before (*N*_0_) and after (*N*) duplicate removal, and recording duration *T*. Human: Lancaster protocol (O*n*); mouse cortical: Paşca protocol (MO*n*).

| Dataset | *N*<sub>0</sub> | *N* | *T* (min) | Dataset | *N*<sub>0</sub> | *N* | *T* (min) |
|---|---|---|---|---|---|---|---|
| O1 | 26 | 25 | 3 | MO5 | 199 | 188 | 10.1 |
| O2 | 131 | 120 | 3 | MO6 | 170 | 146 | 10.4 |
| O3 | 40 | 36 | 3 | MO7 | 234 | 197 | 10.0 |
| O4 | 123 | 122 | 3 | MO8 | 194 | 168 | 10.1 |
| O5 | 151 | 146 | 30 | MO9 | 86 | 78 | 11.0 |
| O6 | 119 | 85 | 30 | MO10 | 157 | 125 | 10.5 |
| MO1 | 215 | 179 | 10.0 | MO11 | 82 | 28 | 12.7 |
| MO2 | 211 | 165 | 10.0 | MO12 | 48 | 44 | 10.0 |
| MO3 | 130 | 113 | 10.1 |  |  |  |  |
| MO4 | 103 | 93 | 10.1 |  |  |  |  |

We compute correlations within short lag windows of 0, 10, and 20 ms (Sec. 3), which probe co-firing at successively longer latencies to allow for conduction and synaptic delays between units; results reported here use the 0 ms window unless noted. Spike trains are smoothed with a 50 ms Gaussian kernel before correlation. This kernel width sets the timescale on which co-firing is measured [17], and 50 ms matches the tens-of-millisecond scale of synaptic interaction among neurons. The activity is strongly bursting, which the null model of Sec. 3 controls for. The sorted units show clear refractory structure, with 0.19% of intervals shorter than the 1.5 ms refractory period, and a pooled median interspike interval of 32 ms (Fig. 1).

**Figure 1.**
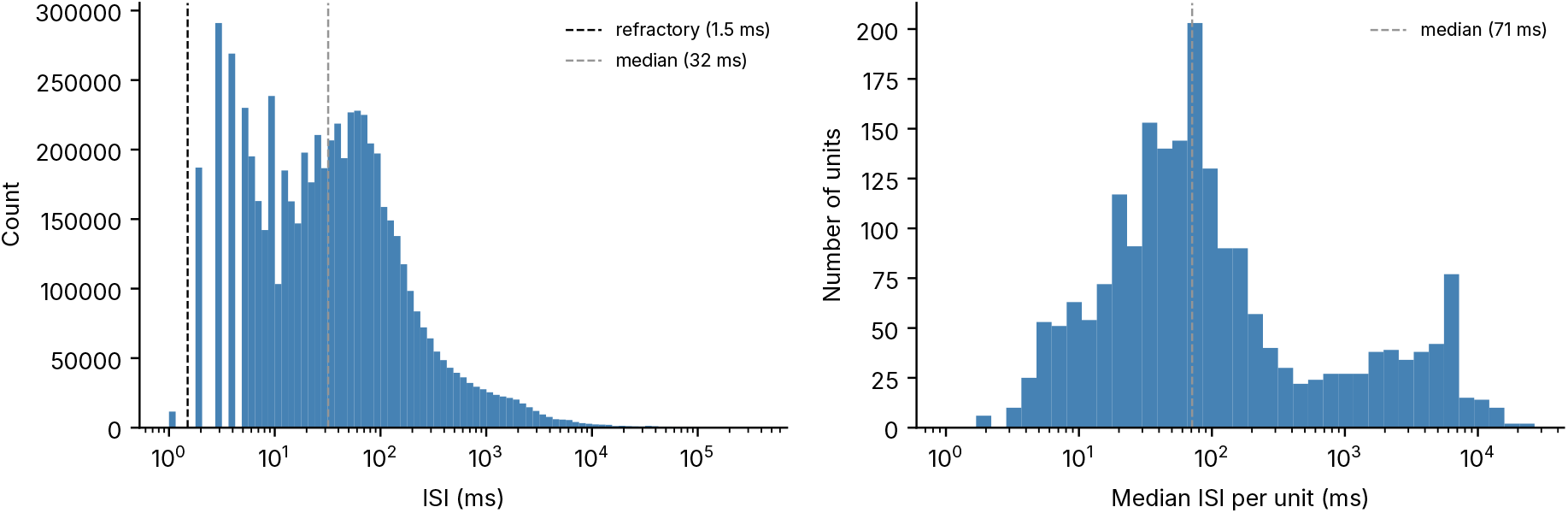
Spike-train statistics across all datasets. (Left) Pooled interspike interval (ISI) distribution over all units and recordings; counts fall to near zero below the 1.5 ms refractory period, confirming well-isolated single units, and the median pooled ISI is 32 ms. (Right) Distribution of per-unit median ISI, whose median across units is 71 ms. Both are consistent with bursting spontaneous activity.

**Figure 2.**
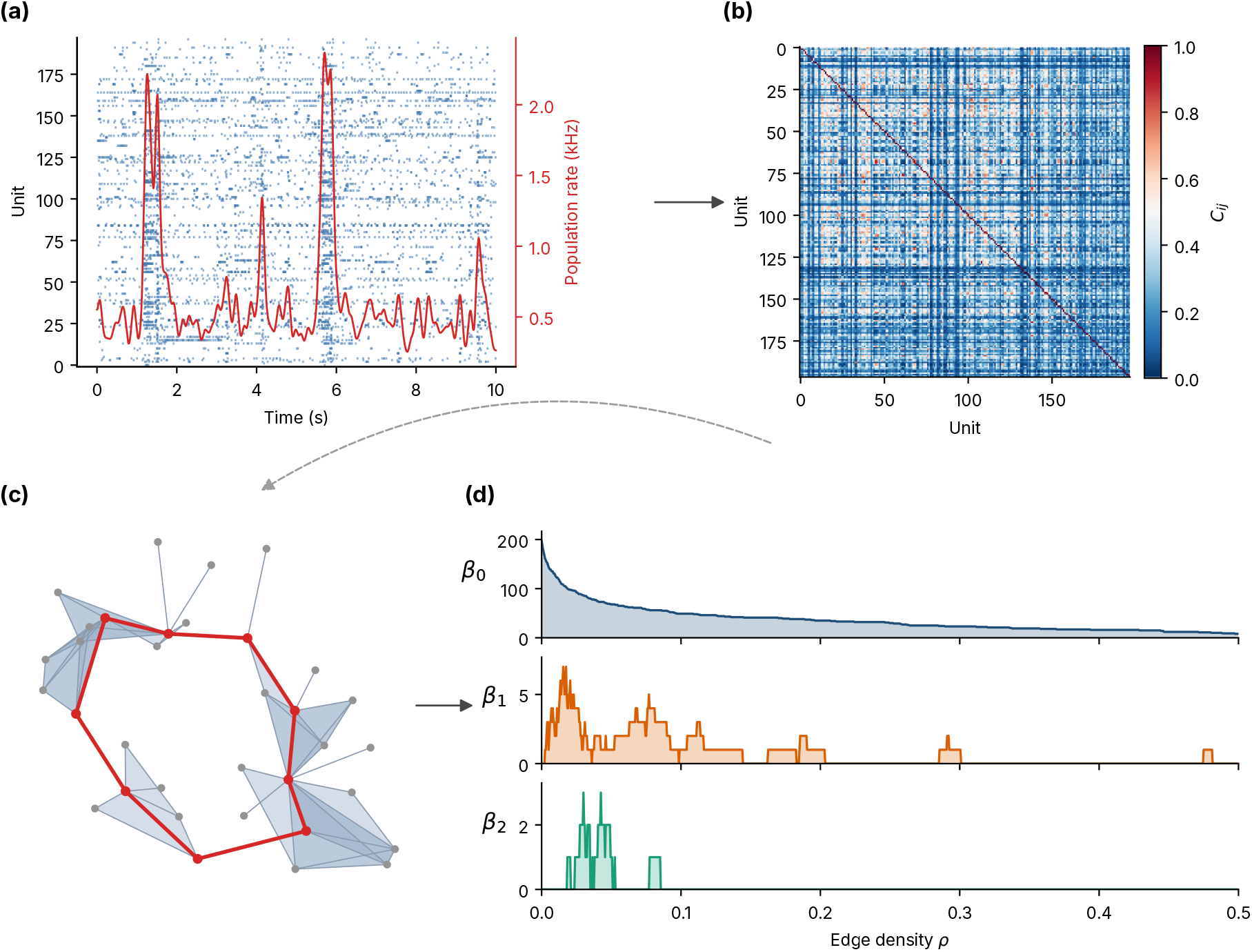
Analysis pipeline, shown for MO7. (a) Spike-sorted units and the population rate. (b) The pairwise correlation matrix *C*_*ij*_. (c) The matrix defines a Vietoris-Rips filtration on *d*_*ij*_ = 1 *− C*_*ij*_, from which persistence reads loops and voids. Shown is a neighborhood of MO7’s most persistent *H*_1_ loop at the scale where it forms, drawn as units (nodes), co-firing edges, and filled 2-simplices (shaded), with the loop highlighted in red enclosing the hole it bounds. Node positions come from multidimensional scaling of the correlation distances, not electrode coordinates. (d) Persistent homology is read out as Betti curves *β*_0_, *β*_1_, *β*_2_ against edge density *ρ*.

Because one neuron can be sorted into several units on a high-density array, we flag a pair as duplicates when more than half of the spikes of the less active unit fall within ±1 ms of a spike of the other, with at least 10 such coincidences and a zero-lag count more than ten times that expected from lags of 4 to 13 ms. We keep the most active unit of each fiagged group, which removes 361 of 2,419 units. A lag-aware variant, which also catches copies offset by up to 5 ms and ignores units with fewer than 100 spikes, removes 228 units. With it the data exceed all 19 surrogates in the datasets that are significant in the main test, and in MO11 as well (Table A1).

## 3. Pairwise time correlations and the null model

For each pair of units (*a, b*) we form a correlation *C*(*a, b*) ∈ [0, 1] from the overlap of their Gaussian-smoothed spike trains,

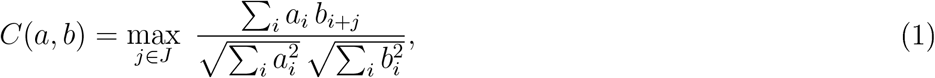

the largest normalized overlap of the two smoothed trains over lags *j* in a window *J* = [*−w, w*], with *C* = 1 for identical activity. At zero lag this is the Gaussian-smoothed spike-train correlation of Schreiber et al. [17]. Built from co-activation, *C* is Hebbian in character, registering units that fire together and underweighting anti-correlated, inhibitory-like coupling.

For *w >* 0 the maximum over lags can only raise *C*, most for pairs whose co-firing is offset in time, and data and surrogates pass through the same operation. We repeated the *H*_1_ test at *w* = 10 and 20 ms against 20 surrogates per dataset. The data exceed all of them in 13 of the 14 datasets significant at zero lag, and integrated *β*_1_ changes by a median of 5-6% in the 13 datasets where it exceeds 0.04. Because *J* is symmetric about zero, *C* is symmetric, as the undirected filtration below requires. Each unit is then a point in a *correlation space* where proximity means functional similarity, not anatomical distance. A correlation links units that co-fire and need not mark a direct synaptic connection: the network is functional, not structural. We test directly whether the resulting topology is intrinsic or inherited from the electrode layout, in the five datasets that retain electrode coordinates (Sec. 5.5).

In the two datasets of Fig. 3 the correlation distributions are right-skewed. Most unit pairs are weakly to moderately correlated, with a peak near *C ≈* 0.1 – 0.2 and a long tail of strongly co-active pairs, and about a tenth of all pairs exceed *C* = 0.6. This tail is a signature of the bursting dynamics in which sub-populations co-activate, reported before in cortical organoids [14, 18, 19], and it motivates a topological treatment: the strong-correlation backbone those pairs form defines a sparse graph whose clique structure is the object of interest.

**Figure 3.**
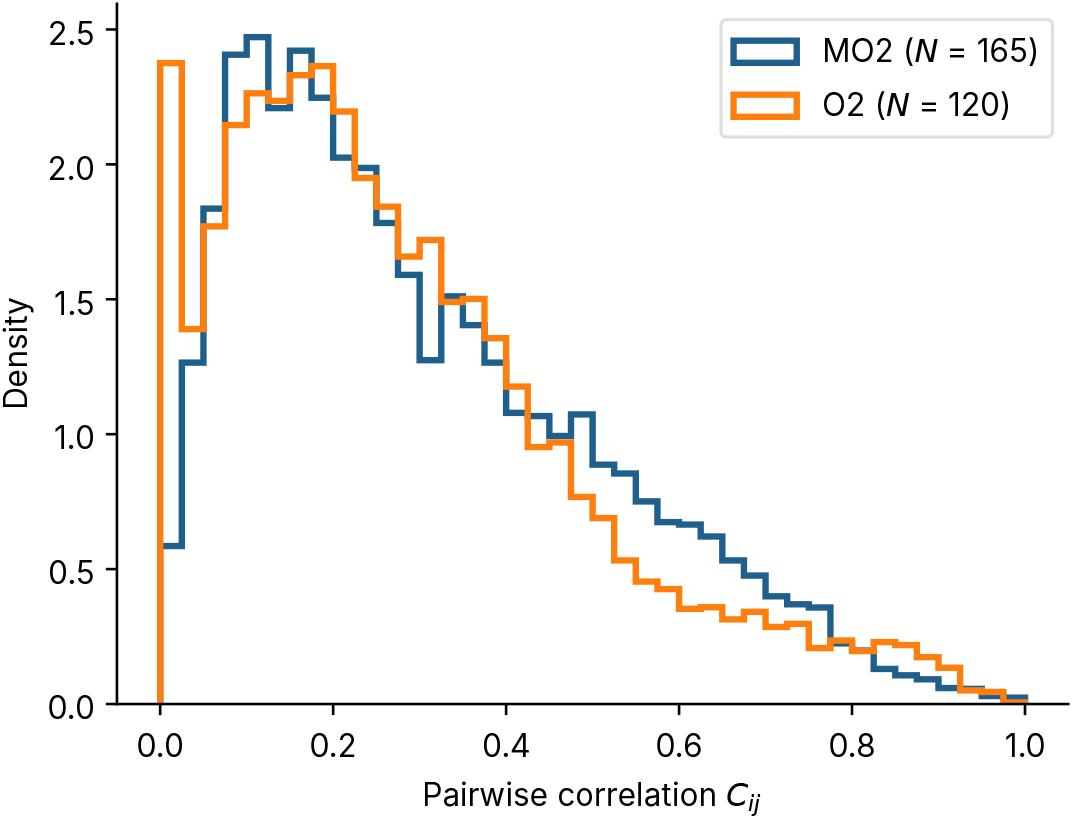
Pairwise correlation distributions for a representative human (O2) and mouse (MO2) dataset: both are right-skewed, with most pairs weakly to moderately correlated and a tail of strongly co-active pairs.

To test whether topological structure exceeds what firing rate and population bursting alone produce, we randomize the binary spike raster under a constraint: the *raster-marginals* model [20]. Each randomization holds fixed both margins of the units×time-bins raster (1 ms bins), every unit’s total spike count and every time bin’s total population activity, while destroying the specific co-firing of particular units, by repeatedly selecting 2 × 2 submatrices of the form 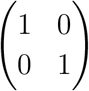 or 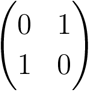 and interchanging the two patterns. Such interchanges connect all 0/1 matrices with given margins 21, but a chain of finite length must also mix [22]. We attempt 10 interchanges per spike, and integrated *β*_1_ and the correlations of the surrogates converge by 2 attempts per spike (Fig. A1). Correlations are recomputed on each randomized raster, and the entire topological analysis is repeated on the surrogate ensemble. Because the surrogates preserve each unit’s firing rate and the population activity in each time bin, the two properties that dominate organoid spiking 20, any topology that exceeds the null reflects co-firing among specific units that these marginals leave undetermined. In the ten- and thirty-minute recordings other than MO11, at most 5 of 100 surrogates contain any loop. Their correlations are close to a function of single-unit properties, since a fit log *C*_*ij*_ = *a*_*i*_ + *a*_*j*_ reproduces their rank order with Spearman coefficient 0.998 in MO7, against 0.84 for the data, and a matrix of exactly that form has no loops at any edge density. The two halves of each recording give correlation matrices that agree with median Pearson *r* = 0.97, and both halves contain loops in every dataset with significant *H*_1_. A fully random (Erdős – Rényi) network serves only as a structural reference, not a significance test.

## 4. Topological methods

Persistent homology represents the correlation network as a growing family of simplicial complexes and counts the holes in them (Fig. 4). The building blocks are simplices: a point, an edge, a filled triangle, and a filled tetrahedron are the simplices of dimension 0 through 3, and a simplicial complex is a collection of them joined along shared faces.

**Figure 4.**
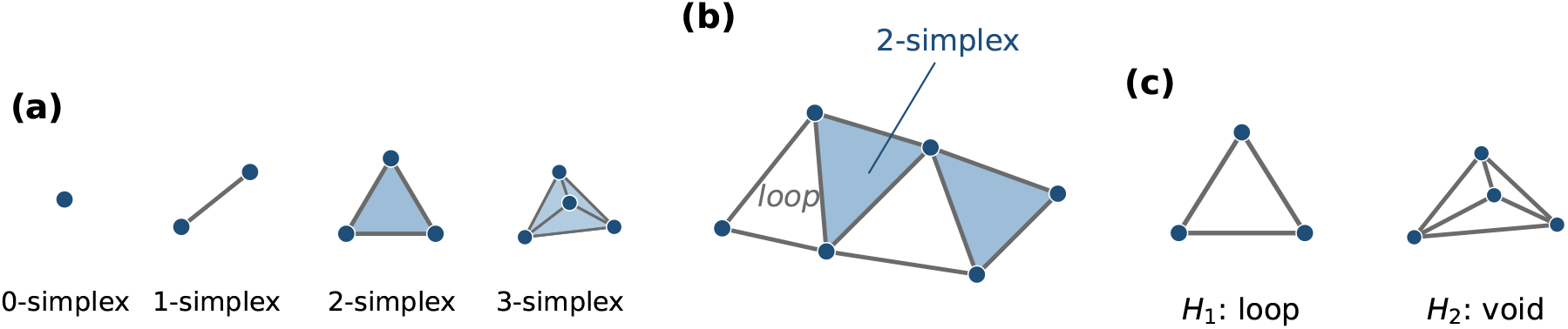
Building blocks of persistent homology. A filled (blue) region is a simplex present in the complex; a region bounded by edges or faces but left unfilled is a hole. (a) Simplices of dimension 0 to 3: a point, an edge, a filled triangle, and a filled tetrahedron. (b) A complex glued from such simplices: the blue triangles are 2-simplices, while the three edges at lower left bound an open triangle that no 2-simplex fills, a loop. (c) The holes the method counts: a loop that no simplex fills is a first-homology (*H*_1_) feature, and an enclosed cavity is a second-homology (*H*_2_) feature.

Persistence is computed on the full correlation matrix with `Ripser` [23] (version 0.6.15, homology up to dimension 2, no distance threshold). We convert each correlation matrix to a dissimilarity *d*_*ij*_ = 1 *− C*_*ij*_ and build the Vietoris – Rips (clique) filtration: as a scale *ε* increases from 0 to 1, an edge appears between units *i* and *j* once *d*_*ij*_ *≤ ε*, and any set of mutually connected units is filled in as a simplex (Fig. 5). The *k*th Betti number *β*_*k*_(*ε*) counts the independent *k*-dimensional homological features present at scale *ε*: *β*_0_ connected components, *β*_1_ loops, *β*_2_ enclosed voids. As *ε* grows, each feature is born at the scale *ε*_birth_ where its cycle first closes and dies at *ε*_death_ where that cycle is filled in.

**Figure 5.**
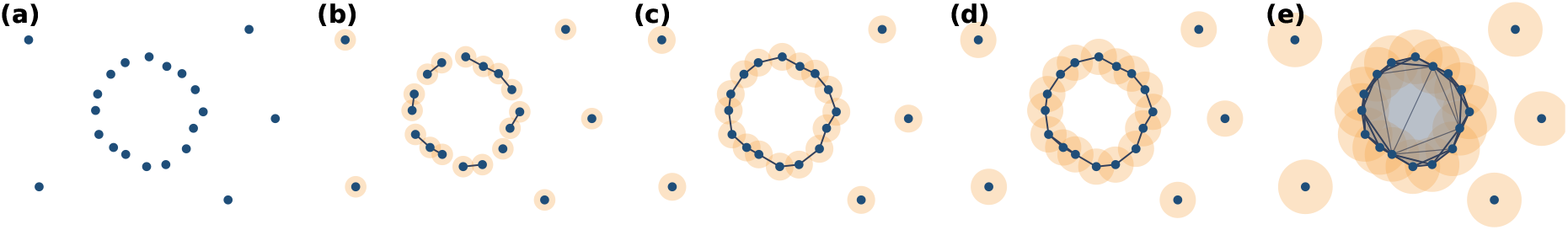
Vietoris-Rips filtration, drawn in two dimensions for intuition; the real filtration runs in the high-dimensional correlation space. Each unit is a point (a), and as the scale *ε* grows, every unit carries a disk of radius *ε/*2; two units join by an edge when their disks overlap (*d*_*ij*_ *≤ ε*), and mutually joined sets fill in as simplices. A loop forms once the ring of edges closes (c), survives as *ε* increases (d), and dies when the interior fills (e). The range of *ε* over which it survives is its persistence, which separates a real feature from noise.

Rather than fix a single scale, we report *density*-*indexed* Betti curves, the clique-topology construction introduced for neural correlation matrices by Giusti et al. 4. The edge density *ρ ∈* [0, 1] is the fraction of the ^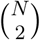^ possible pairs present at a given scale; it is a monotone reparametrization of *ε*. Writing *F* for the empirical cumulative distribution function of the pairwise correlations, *ρ* = 1 *− F* (*θ*) at correlation threshold *θ* = 1 *− ε*, the fraction of pairs whose correlation exceeds *θ*. Plotting *β*_*k*_(*ρ*) and reporting the integrated Betti value ∫ *β*_*k*_(*ρ*) *dρ* has two virtues: it does not depend on the particular correlation scale, and it places data and surrogate on a common, rank-matched axis, so the null comparison is made at equal sparsity even though the surrogates’ correlation values differ in scale. We use the undirected construction throughout; directed (flag) complexes are an extension to directed data. A separate threshold at the 50th density percentile is used only for the supporting graph-theoretic measures. Each loop or void is a point (*ε*_birth_, *ε*_death_) in a persistence diagram. The bottleneck distance between two diagrams is the smallest *δ* for which every point of one can be matched to a point of the other, or to the diagonal, within *δ*. It measures how much the topology changes between two conditions, and we use it in Sec. 5.2.

A Vietoris – Rips complex can have homology in arbitrarily high degree even for points in the plane [24], so the homological dimension of a feature does not reflect the spatial dimension of the recording.

## 5. Results

This section reports five results. First, organoid networks carry more loop (*H*_1_) structure than the rate- and population-preserving null produces (Sec. 5.1). Second, this structure rests on a non-redundant core of strongly co-active units (Sec. 5.2). Third, the number of homological dimensions a network populates grows with its size (Sec. 5.3). Fourth, enclosed voids (*H*_2_) emerge only in the larger networks (Sec. 5.4). Finally, the loop structure reflects co-firing rather than the physical layout of the array (Sec. 5.5).

### 5.1. Loop structure exceeds the rate- and population-preserving null

Our primary statistic is the integrated first Betti number [4], *β*_1_(*ρ*) *dρ*, the area under the density-swept loop count. It summarizes, in a single threshold-free number, how much loop structure a network carries across the whole filtration. Because it is indexed by edge density rather than by raw correlation, it can be compared directly between a recording and its surrogates even though randomization shifts the correlation values (Sec. 4). For each dataset we compare this quantity to the distribution obtained from 100 raster-marginals surrogates. The null distributions are not normal (Fig. A2), so we report the rank-based *p*-value *p* = (*r*+1)/101, where *r* is the number of surrogates whose integrated *β*_1_ is at least that of the data [25], together with *q*, the *p*-value corrected for testing 18 datasets [26].

Loop structure in the organoid data exceeds the surrogate null in 14 of the 18 datasets, all with *q ≤* 0.025 (Fig. 6a, Table A1). The four that do not separate from the null are the two smallest networks (O1, *N* = 25; MO11, 28) and two of the larger human sets (O4, 122; O2, 120). Size alone does not decide the outcome. O1 has no loops, and O2 and O4 are three-minute recordings whose surrogates also contain loops. Where the null is nearly empty, a dataset with any loops is significant. In the mid-to-large networks integrated *β*_1_ ranges from 0.045 to 0.44.

**Figure 6.**
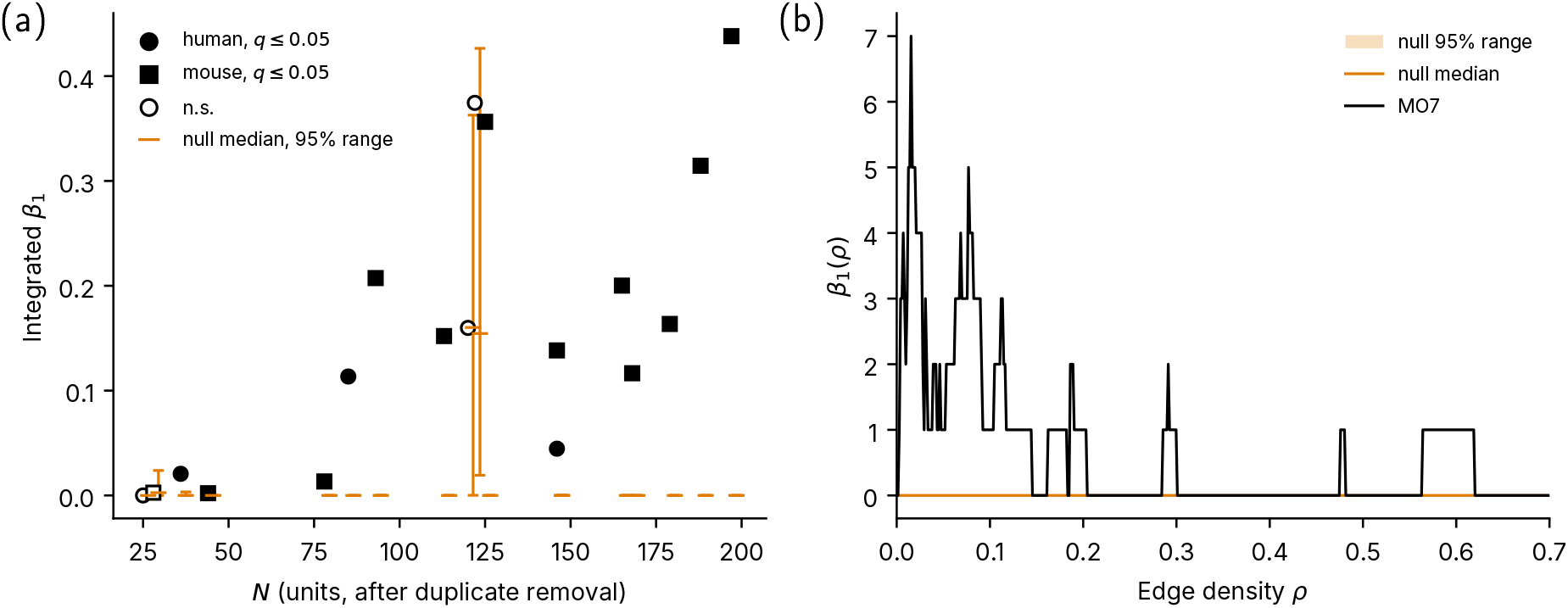
(a) Integrated *β*_1_ for data (filled, *q ≤* 0.05; open, n.s.; circles, human; squares, mouse) against the raster-marginals null (median and central 95%) across all 18 datasets, versus *N*. (b) *β*_1_(*ρ*) for MO7: data (black) versus null median and central 95% (orange).

Most loops occur at low edge density. In MO7, *β*_1_(*ρ*) peaks near *ρ* = 0.016 and at most three loops remain beyond *ρ* = 0.1 (Fig. 6b). As weaker correlations enter the filtration, the loops are filled in by the triangles that high clustering supplies. A few loops persist to *ρ ≈* 0.6. The topology is therefore mainly a property of the strong-correlation backbone: the loops are carried by the strongly co-active units, the “choristers’ of the population rather than the weakly coupled soloists 27. Because the surrogates hold each unit’s firing rate and each time bin’s population activity fixed, the loop structure that exceeds them cannot be a by-product of rate or of the population-wide bursts that dominate organoid activity. It reflects co-firing structure beyond the reach of those first-order properties.

### 5.2. The loops occupy a non-redundant core

If the loop structure were diffuse, supported redundantly by many interchangeable units, it would degrade gracefully as units are removed. If instead it rests on a specific set of carrying units, removing those should disrupt it sharply while removing others should barely matter. We test this by deleting 10% of the units and recomputing the topology, contrasting random deletion with deletion drawn uniformly from the supports of the *H*_1_ cocycle representatives (14-44% of the units), each over 100 draws of the same number of units.

Random removal leaves the loop structure largely intact: across the datasets with loop structure to test, integrated *β*_1_ is retained at a median of 94%, against 71% under targeted removal (Fig. 7a), so no small, interchangeable subset is doing all the work. Targeted removal disrupts it far more, but quantifying that disruption requires care. The natural-seeming measure, the change in loop count, is in fact misleading: deleting a densely connected hub sparsifies its neighborhood and can new loops, so *β*_1_ sometimes rises under targeted removal even as the original loops are destroyed. We therefore measure disruption by the bottleneck distance between the original and post-removal *H*_1_ persistence diagrams, which registers how far the loop structure has moved regardless of whether new features appear. The mean bottleneck distance under targeted removal exceeds that under random removal, a ratio above 1, in all 14 datasets with enough loop structure to compare. We set aside the four lowest-count sets (O1, MO11, O3, MO12), whose cocycle supports hold fewer units than are deleted. The median ratio is 1.54, with a range of 1.21 to 1.66 (Fig. 7b, Table A1), and a ranked test across datasets gives one-sided *p* = 6.1 × 10^*−*5^ (Wilcoxon). Within each dataset the 100 targeted draws exceed the 100 random draws (one-sided rank-sum test, *p ≤* 2.3 × 10^*−*5^ in all 14). Loop-supporting units are more strongly connected than the rest (median ratio of weighted degree 1.23), so for each targeted draw we also delete the units outside the supports closest in weighted degree. Targeted removal moves the *H*_1_ diagram more than this strength-matched removal in 13 of the 14 datasets, and about as much in MO8 (median ratio 2.53, *p* = 6.1 × 10^*−*5^). The loops are thus carried by an identifiable, non-redundant core of units rather than spread interchangeably across the population.

**Figure 7.**
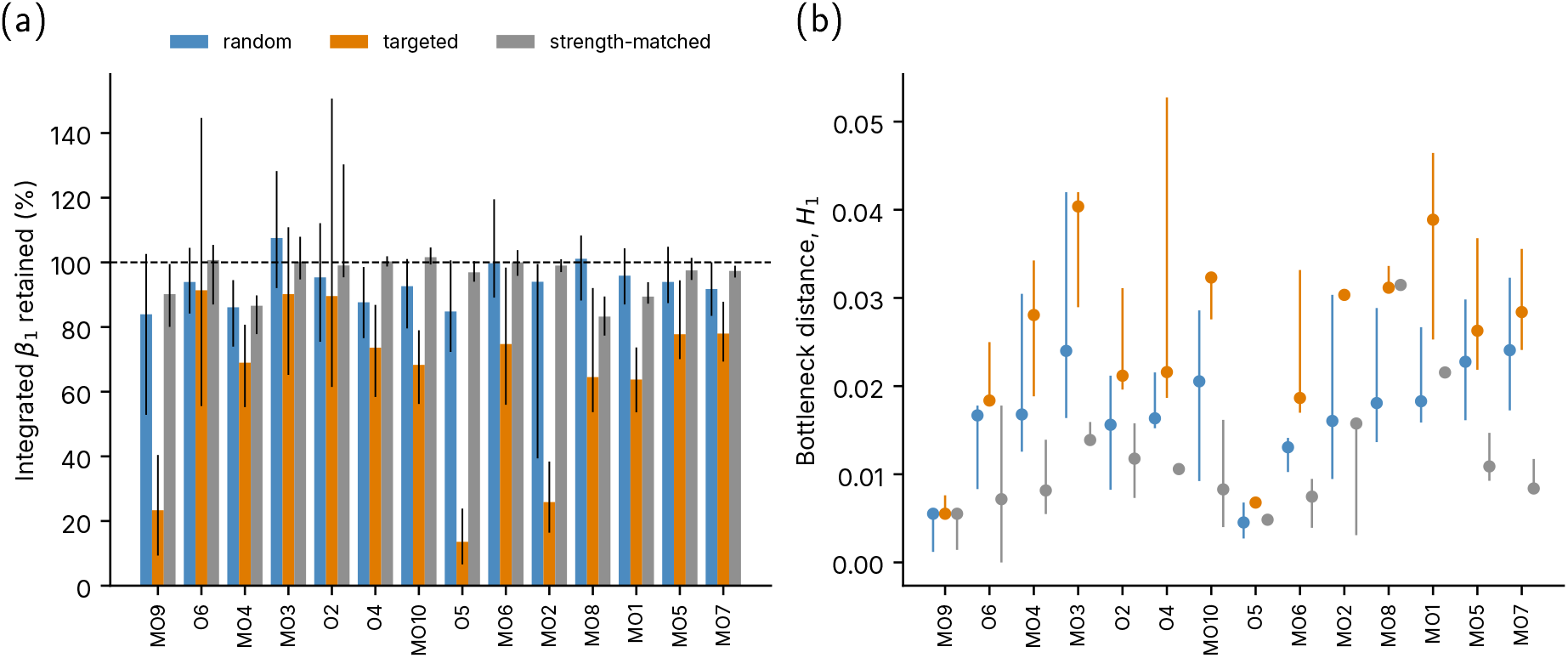
(a) Fraction of integrated *β*_1_ retained under random (blue), targeted (orange), and strength-matched (grey) removal of 10% of units, median and interquartile range over 100 draws; dashed line: full retention. (b) Bottleneck distance of the *H*_1_ diagram under the same three conditions.

### 5.3. Topological richness grows with network size

Richness here means how many homological dimensions a network resolves above the null, that is, how many of *H*_0_, *H*_1_, *H*_2_ it populates. It climbs with *N* because each Betti number becomes resolvable above the null only as the network grows, with successive dimensions requiring more units. The transition is statistical, not a sharp threshold. Connected components (*H*_0_) resolve at every size. Loops (*H*_1_) need enough units to close a cycle and resolve above the null in 14 of 18 datasets, not in strict order of size (Sec. 5.1). Enclosed voids (*H*_2_) come online later, clearing the null only among the larger networks (Sec. 5.4). A larger network is thus more likely to resolve each next dimension: *H*_1_ appears readily and *H*_2_ in the larger networks. The count within a dimension tells the same story more weakly: integrated *β*_1_ rises with *N* (Pearson *r* = 0.68; Fig. 8), but the correlation is inflated by the small networks that resolve no loops, so the within-dimension magnitude is not a reliable size law. Richness grows with size by adding dimensions, not loops within one.

**Figure 8.**
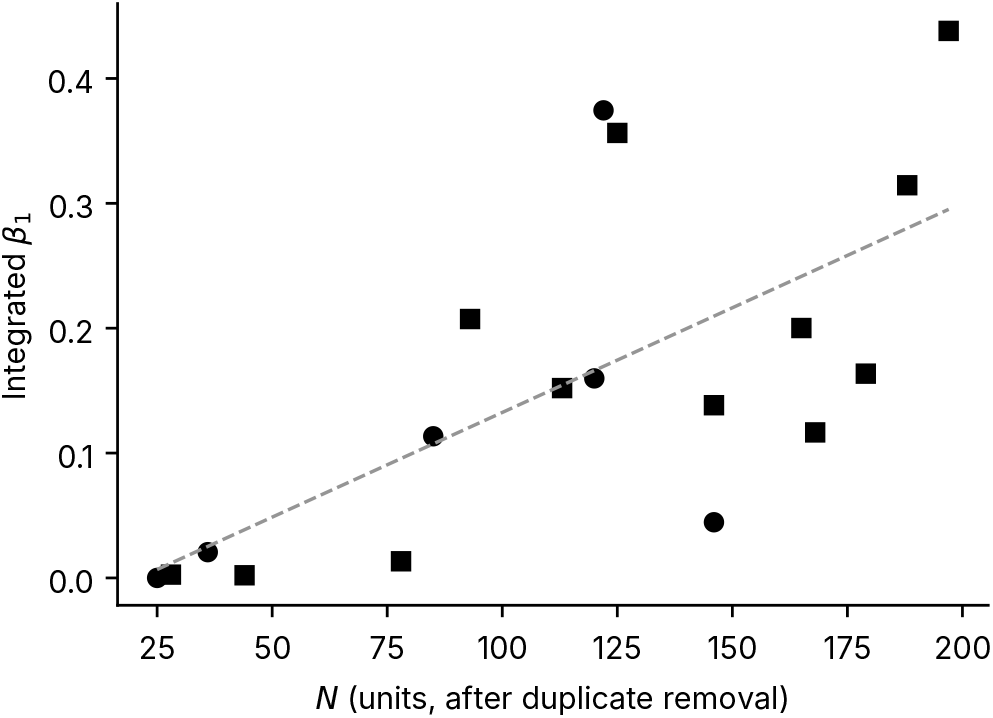
Integrated *β*_1_ against the number of units *N* (circles, human; squares, mouse; dashed line: linear fit, *r* = 0.68).

Random subsets of MO7 exceed their own null in *H*_1_ at every size from 25 units. In *H*_2_ they exceed it in none of 10 subsets of 25 units, in 7 of 10 subsets of 50, and in all subsets of 100 and 150 (Fig. A3). Within a recording voids thus need more units than loops, as across datasets. Our networks are at the scale of the 88 place cells of Giusti et al. [4] and two orders of magnitude smaller than the reconstructed microcircuit of Reimann et al. [6].

### 5.4. Second homology appears in the larger networks

Enclosed voids (*H*_2_) require both more units and richer co-firing to form than loops. Under the same surrogate test, *H*_2_ rises significantly above the null in nine datasets (O6, MO3, MO10, O5, MO6, MO8, MO1, MO5, MO7), each with *N ≥* 85 and *q* = 0.020, and most strongly in the largest (Fig. 9a; MO7 in Fig. 9b). No dataset below *N* = 85 shows significant *H*_2_, but neither do four of the thirteen at or above it (MO4, O2, O4, MO2), so this size is necessary in this corpus but not sufficient. The voids that drive it are few and low-persistence, shorter-lived than the longest loops in the same data, so the excess is significant but not yet a robust feature.

**Figure 9.**
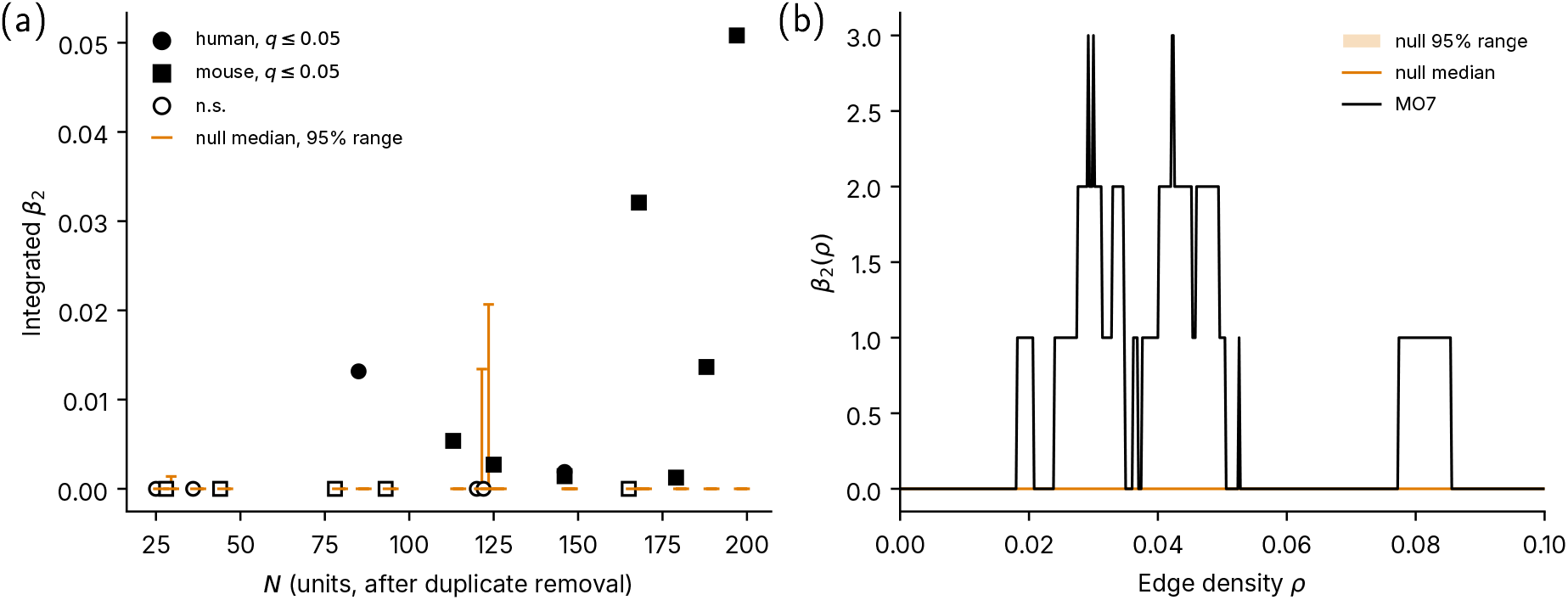
(a) Integrated *β*_2_ for each dataset against unit count *N*. Filled symbols mark datasets whose *β*_2_ exceeds the rate- and population-preserving null at *q ≤* 0.05, open symbols those that do not (circles, human; squares, mouse). Orange marks give the null median and central 95%. (b) *β*_2_(*ρ*) for MO7: data (black) and null (orange).

### 5.5. The loops reflect co-firing, not the electrode layout

A correlation network could in principle report the geometry of the recording rather than the organization of the activity, and for the five datasets that retain electrode coordinates (O2-O6) we can check. Correlation falls with electrode distance in four of them (Spearman *ρ* = *−*0.10 to *−*0.71 over all unit pairs) and not in O5 (0.01) (Fig. 10c).

**Figure 10.**
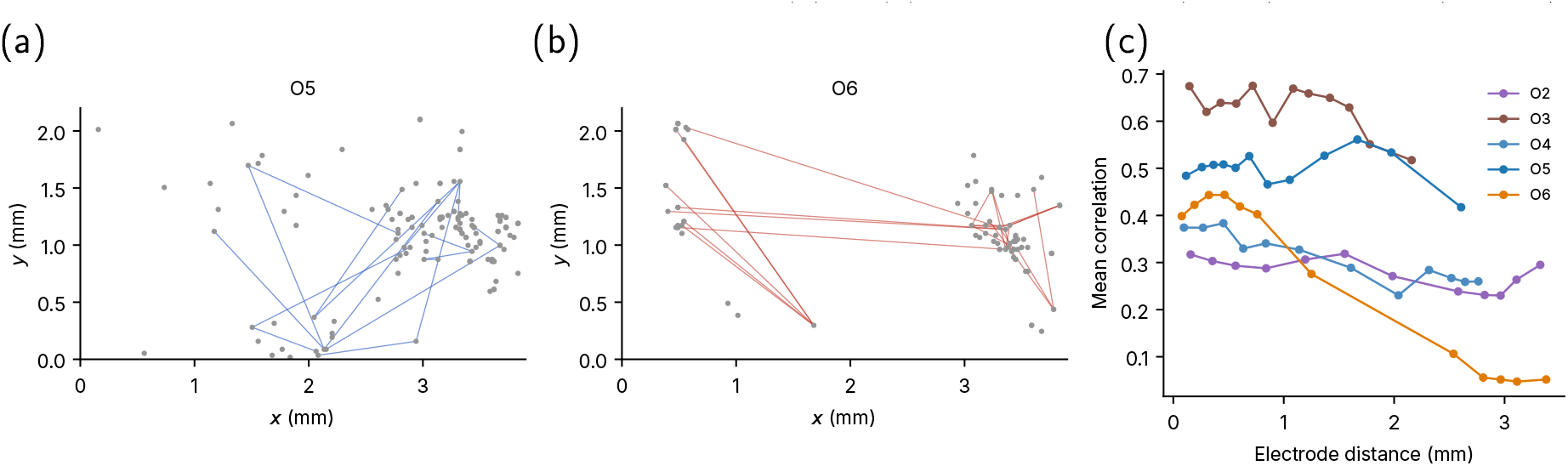
Where the loops occur. (a,b) Unit positions on the array (mm, *x* along its long axis) with the edges of the *H*_1_ cocycle supports overlaid, for O5 (a) and O6 (b). In both, the supports span the array. (c) Mean correlation versus electrode distance for O2-O6, in bins of equal pair count.

If the loops were inherited from the layout, the units in their *H*_1_ cocycle supports would cluster on the array. In O2, O4, O5 and O6 their mean electrode distance is 0.96 to 1.13 times that of random unit sets of the same size, and none is more compact than random sets (*p ≥* 0.20), not even O6, whose correlations depend strongly on distance (Fig. 10b). In O3, whose support holds only three units, the ratio is 0.40 (*p* = 0.06).

## 6. Conclusion

Persistent homology resolves real, structured loop topology in spontaneous organoid activity at the modest node counts that recordings provide. The loop structure exceeds a null that matches firing rate and population bursting, concentrates in a non-redundant core of carrying units, and is joined by second homology (*H*_2_) in the larger networks. Neither the electrode layout, in the five datasets with coordinates, nor the lag window, in 13 of the 14 significant datasets, accounts for this structure.

Topological analysis of neural systems has been held back by the suspicion that meaningful Betti numbers require unattainably many nodes. Our results show otherwise. Across eighteen datasets, *H*_1_ structure is resolvable above the null in 9 of the 11 datasets with *N ≥* 100 and in five of the seven smaller ones. This shows persistent homology to be a usable instrument for the recordings the field already has, and it points to a clear next step: applying the same pipeline to optical recordings with single-cell resolution and, ultimately, to three-dimensional recording at high temporal resolution. Because a planar array captures only a fraction of the units in a three-dimensional culture, raising the unit count this way should bring out more of the second homology that appears here. A directed filtration would add the firing order the undirected construction sets aside, and applying the same null test across the two protocols would show whether the carrying core is conserved across them.

## Acknowledgements

We thank Ginestra Bianconi and Michael Freedman for helpful insight into multiple aspects of simplicial complexes and topological data analysis. We acknowledge early experimental support and discussions with Raymond Griffard and Tjitse van der Molen, and theoretical support via discussions with Om Biyani and Bismah Rizwan. We thank Bismah Rizwan for a close reading of the analysis code. This work was performed in part with support by the Google Academic Research Award program (LDC, KSK, DB) and the U.S. National Science Foundation under Grants DGE-2125899 (MB, LDC) and PHY-2515059 (LDC). NM gratefully acknowledges funding from the Noyce Foundation. LDC thanks the Institute of Science and Technology Austria for hospitality, and Maksym Serbyn for hosting him during the sabbatical in which this work was revised and completed.

## Data availability

The data and analysis code that support the findings of this study are available upon request from the authors.

## CRediT

Conceptualization (all); Data curation (EB, MB); Formal analysis (EB, MB, LDC); Funding acquisition (LDC, KSK); Investigation (EB, SH, LF); Methodology (EB, MB, NM, LDC); Project administration (LDC, DB, KSK); Resources (KSK); Software (EB, MB, LDC); Supervision (LDC, DB, KSK); Validation (EB, MB, LDC); Visualization (all); Writing - original draft (EB, LDC, DB); Writing - review & editing (all)

## Appendix A Per-dataset statistics and null diagnostics

This appendix collects the per-dataset results and the checks on the surrogate null. Table A1 lists for each dataset the integrated Betti numbers 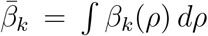 of the data, the median and 97.5th percentile of the null, *r* and *q* (Sec. 5.1). For the lag-aware duplicate criterion it gives the unit counts and *r* out of 19 surrogates. The last two columns are the removal ratios of Sec. 5.2, targeted over random (T/R) and targeted over strength-matched (T/S).

Fig. A1, computed before duplicate removal, shows that the surrogate chain converges by 2 attempts per spike, well below the 10 used here, while a fixed 10^5^ attempts (dashed lines) leaves MO7 and O6 unconverged. Fig. A2 shows the null distributions of O2 and O4, the only datasets above 50 units whose null is not concentrated at zero. Both are bounded at zero and skewed. Fig. A3 shows the subsampling of MO7 described in Sec. 5.3.

**Figure A1.**
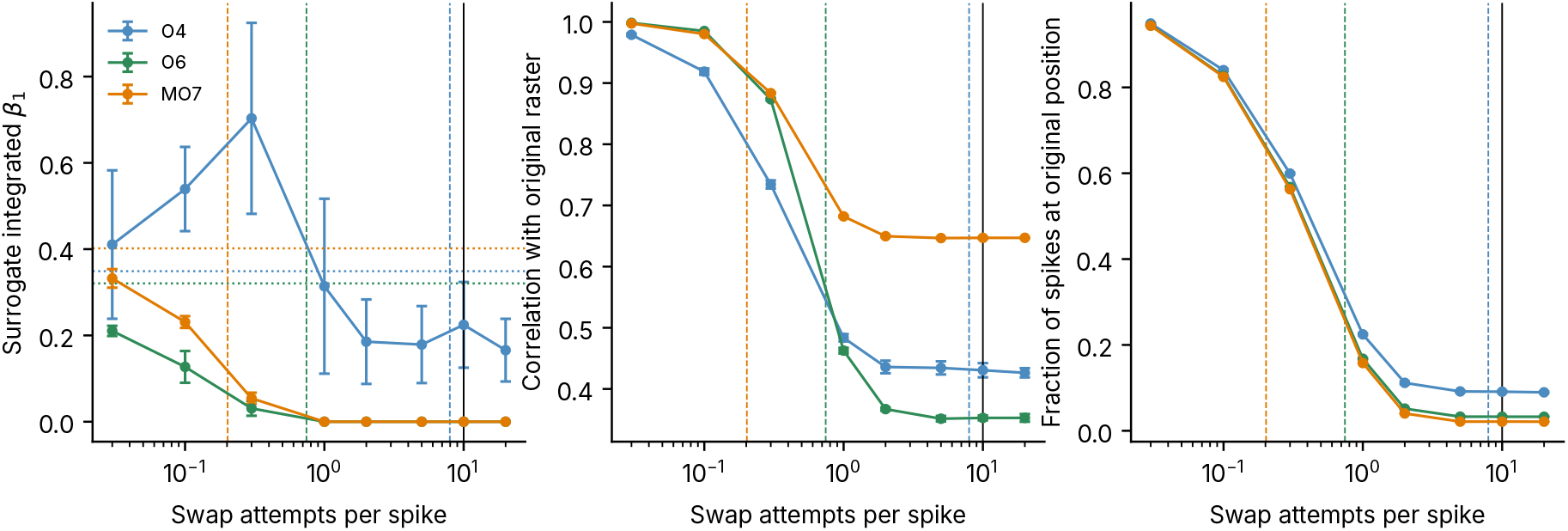
Convergence of the raster-marginals chain (mean *±* sd). (Left) Integrated *β*_1_ of the surrogates, data dotted. (Centre) Correlation of surrogate with data matrix. (Right) Fraction of spikes at their original position. Dashed: 10^5^ attempts; solid: 10 attempts per spike.

**Figure A2.**
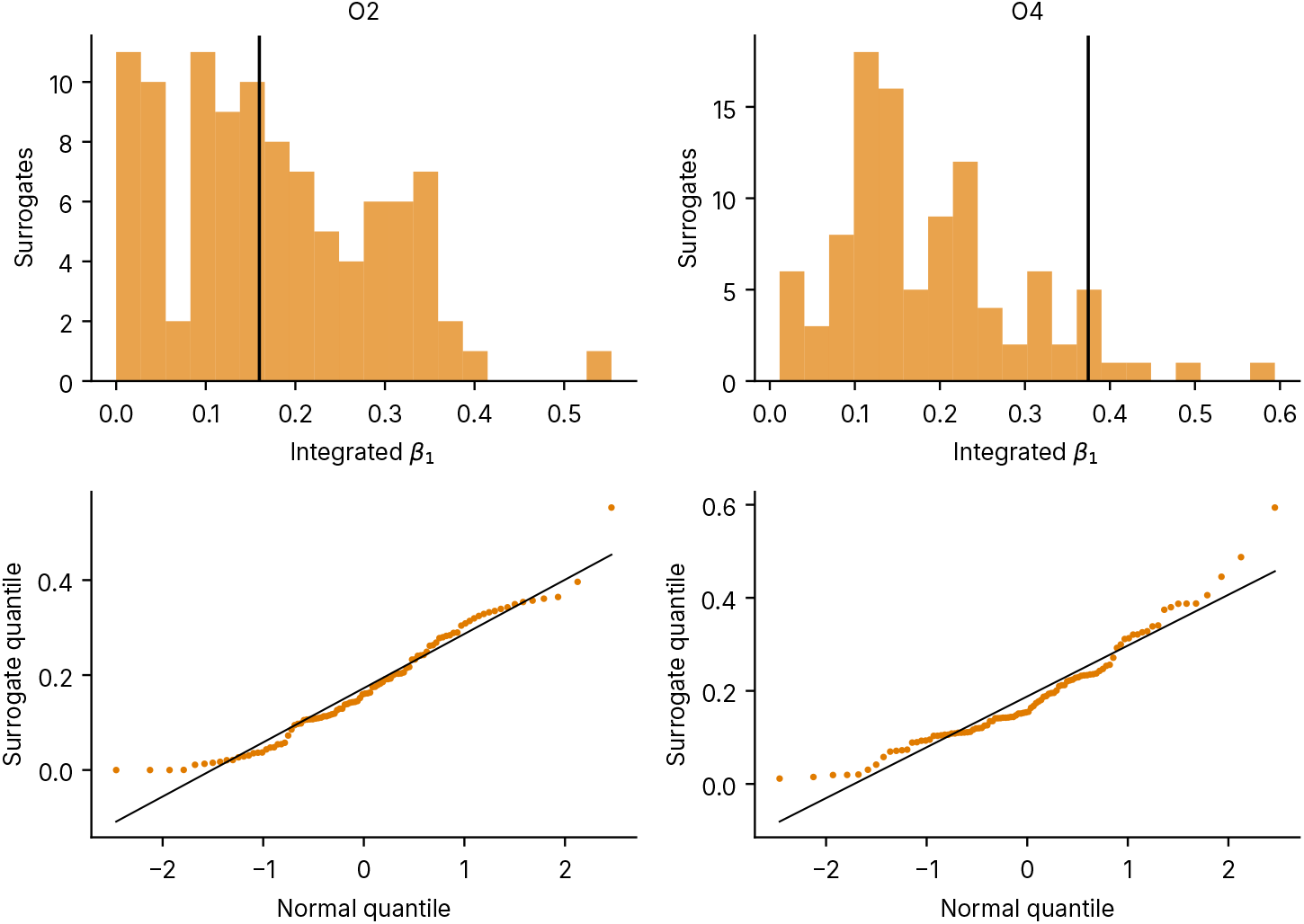
Null distributions of integrated *β*_1_ for O2 and O4 (data as vertical line), with normal QQ plots below.

**Figure A3.**
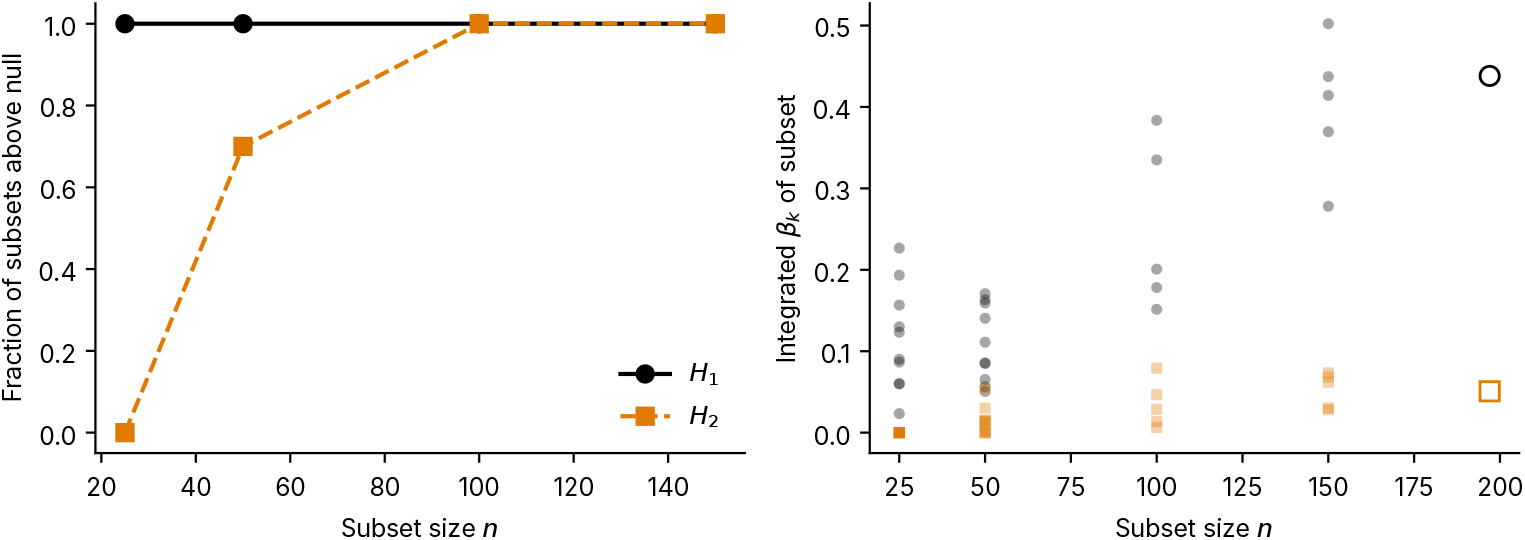
Subsampling MO7. (Left) Fraction of subsets that exceed their own null in *β*_1_ (black) and *β*_2_ (orange), against subset size *n*. (Right) Integrated *β*_1_ (grey) and *β*_2_ (orange) of each subset; open symbols, full network.

**Table A1.**
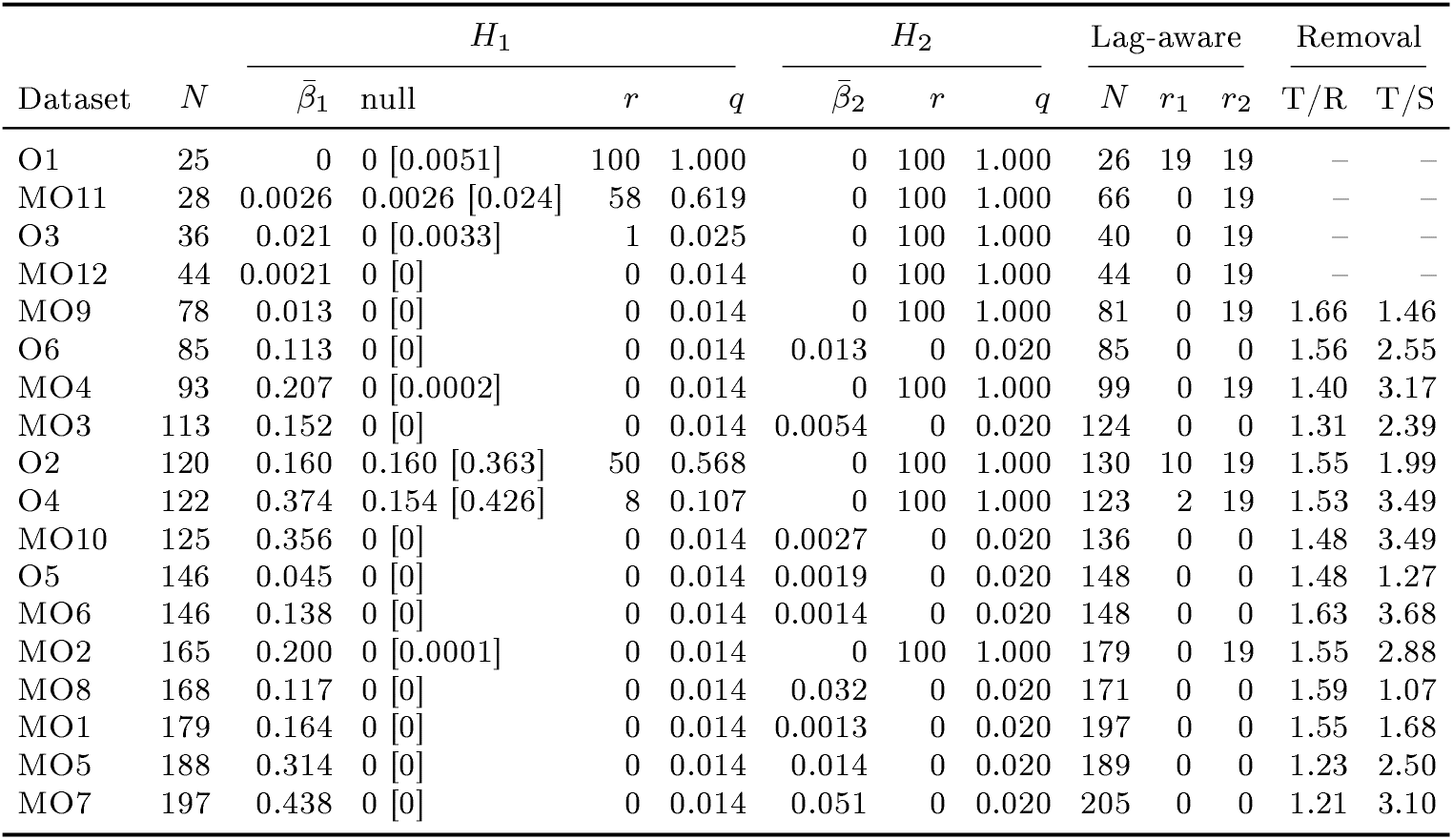
Per-dataset statistics, ordered by *N*. Null: median [97.5th percentile].

|  |  | *H*<sub>1</sub> |  |  |  | *H*<sub>2</sub> |  |  | Lag-aware |  |  | Removal |  |
|---|---|---|---|---|---|---|---|---|---|---|---|---|---|
| Dataset | *N* | *β̄*<sub>1</sub> | null | *r* | *q* | *β̄*<sub>2</sub> | *r* | *q* | *N* | *r*<sub>1</sub> | *r*<sub>2</sub> | T/R | T/S |
| O1 | 25 | 0 | 0 [0.0051] | 100 | 1.000 | 0 | 100 | 1.000 | 26 | 19 | 19 | – | – |
| MO11 | 28 | 0.0026 | 0.0026 [0.024] | 58 | 0.619 | 0 | 100 | 1.000 | 66 | 0 | 19 | – | – |
| O3 | 36 | 0.021 | 0 [0.0033] | 1 | 0.025 | 0 | 100 | 1.000 | 40 | 0 | 19 | – | – |
| MO12 | 44 | 0.0021 | 0 [0] | 0 | 0.014 | 0 | 100 | 1.000 | 44 | 0 | 19 | – | – |
| MO9 | 78 | 0.013 | 0 [0] | 0 | 0.014 | 0 | 100 | 1.000 | 81 | 0 | 19 | 1.66 | 1.46 |
| O6 | 85 | 0.113 | 0 [0] | 0 | 0.014 | 0.013 | 0 | 0.020 | 85 | 0 | 0 | 1.56 | 2.55 |
| MO4 | 93 | 0.207 | 0 [0.0002] | 0 | 0.014 | 0 | 100 | 1.000 | 99 | 0 | 19 | 1.40 | 3.17 |
| MO3 | 113 | 0.152 | 0 [0] | 0 | 0.014 | 0.0054 | 0 | 0.020 | 124 | 0 | 0 | 1.31 | 2.39 |
| O2 | 120 | 0.160 | 0.160 [0.363] | 50 | 0.568 | 0 | 100 | 1.000 | 130 | 10 | 19 | 1.55 | 1.99 |
| O4 | 122 | 0.374 | 0.154 [0.426] | 8 | 0.107 | 0 | 100 | 1.000 | 123 | 2 | 19 | 1.53 | 3.49 |
| MO10 | 125 | 0.356 | 0 [0] | 0 | 0.014 | 0.0027 | 0 | 0.020 | 136 | 0 | 0 | 1.48 | 3.49 |
| O5 | 146 | 0.045 | 0 [0] | 0 | 0.014 | 0.0019 | 0 | 0.020 | 148 | 0 | 0 | 1.48 | 1.27 |
| MO6 | 146 | 0.138 | 0 [0] | 0 | 0.014 | 0.0014 | 0 | 0.020 | 148 | 0 | 0 | 1.63 | 3.68 |
| MO2 | 165 | 0.200 | 0 [0.0001] | 0 | 0.014 | 0 | 100 | 1.000 | 179 | 0 | 19 | 1.55 | 2.88 |
| MO8 | 168 | 0.117 | 0 [0] | 0 | 0.014 | 0.032 | 0 | 0.020 | 171 | 0 | 0 | 1.59 | 1.07 |
| MO1 | 179 | 0.164 | 0 [0] | 0 | 0.014 | 0.0013 | 0 | 0.020 | 197 | 0 | 0 | 1.55 | 1.68 |
| MO5 | 188 | 0.314 | 0 [0] | 0 | 0.014 | 0.014 | 0 | 0.020 | 189 | 0 | 0 | 1.23 | 2.50 |
| MO7 | 197 | 0.438 | 0 [0] | 0 | 0.014 | 0.051 | 0 | 0.020 | 205 | 0 | 0 | 1.21 | 3.10 |

## Notes

### Competing Interest Statement

The authors have declared no competing interest.

### Summary of Updates

Revised in response to peer review: duplicate units removed and all results recomputed, null model and statistics updated, new control analyses and an appendix of per-dataset statistics added, figures revised, author affiliations updated.

